# Multiplexed relative and absolute quantitative immunopeptidomics reveals MHC I repertoire alterations induced by CDK4/6 inhibition

**DOI:** 10.1101/2020.03.03.968750

**Authors:** Lauren E Stopfer, Joshua M Mesfin, Brian A Joughin, Douglas A Lauffenburger, Forest M White

## Abstract

Peptides bound to class I major histocompatibility complexes (MHC) play a critical role in immune cell recognition and can trigger an antitumor immune response in cancer. Surface MHC levels can be modulated by anticancer agents, altering immunity. However, understanding the peptide repertoire’s response to treatment remains challenging and is limited by quantitative mass spectrometry-based strategies lacking robust normalization controls. We describe a novel approach that leverages recombinant heavy isotope-coded peptide MHCs (hipMHCs) and multiplex isotope tagging to quantify peptide repertoire alterations using low sample input. HipMHCs improve quantitative accuracy of peptide repertoire changes by normalizing for variation across analyses and enable absolute quantification using internal calibrants to determine copies per cell of MHC antigens, which can inform immunotherapy design. Applying this platform in melanoma to profile the immunopeptidome response to CDK4/6 inhibition and interferon gamma, known modulators of antigen presentation, we uncovered treatment-specific alterations, connecting the intracellular response to extracellular immune presentation.

## INTRODUCTION

Cells present signals on the extracellular surface that serve as targets for immune cell recognition. These signals, peptides presented by class I MHCs, are typically derived from intracellular source proteins, and may therefore provide an external representation of the internal cell state. As a reflection of this, the peptide MHC (pMHC) repertoire, or “immunopeptidome”, of cancer cells may contain tumor-associated or mutation-containing antigens that serve as tumor-specific markers to activate T cells and initiate an anti-tumor immune response. This interaction can be strengthened with checkpoint blockade (CB) immunotherapies, however low response rates and toxicity remain barriers to their broad clinical success.^1,2^ A growing body of evidence suggests that combining CB with other treatments such as small molecule inhibitors, cytotoxic agents, and radiotherapy could potentiate the response to CB, in part by augmenting tumor immunogenicity through increased surface pMHC expression.^3–9^ While clinical trials in this space have shown promise^10,11^, the optimal combination of agents, as well as the order and timing of administration, are only beginning to be understood. In order to improve combinatorial strategies, a quantitative, molecular understanding of how different perturbations shift the immunopeptidome is required. Furthermore, achieving absolute quantification of presented antigens is necessary to inform immunotherapy drug design, as targeted strategies have varying thresholds of antigen expression required for an optimal antitumor response.

Traditional methods to profile pMHC repertoires using mass spectrometry (MS) are well-documented^12–14^ but quantitative methods have critical limitations. Specifically, most common relative quantification pMHC methods lack a normalization strategy to account for variations in sample input and processing.^15–20^ Peptide losses during processing are estimated to be as much as 99.5% but vary across peptides and samples, underscoring the need for normalization.^21^ Absolute quantification of pMHCs to date is most commonly performed by comparing endogenous levels of pMHCs to exogenous peptide standards, again failing to account for sample losses.^22–25^ Losses can be accounted for with internal pMHC standards, but require laborious refolding of pMHCs for every target of interest.^21^ Nevertheless this approach relies on single point calibration, ignoring the effects of ion suppression, thereby inaccurately estimating absolute pMHC levels in quantitative analyses.

To combat these challenges in quantitative immunopeptidomic profiling, we present a platform that utilizes ultraviolet (UV)-mediated peptide exchange of recombinant MHC monomers to generate on demand heavy isotope-labeled pMHCs for relative and absolute quantification of pMHC repertoires using low sample input. We demonstrate that the addition of heavy isotope pMHCs (hipMHCs) spiked into sample lysates for normalization improves quantitative accuracy between samples for both label-free and multiplexed (TMT-labeled) analyses and provides an estimate of ion suppression through regression against a titrated internal calibrant. Furthermore, we utilize hipMHC multipoint embedded standard curves coupled with isobaric mass tags to accurately quantify the absolute number of copies per cell of target antigens within a single analysis. We applied this platform to profile immunopeptidomic changes in melanoma cell lines, comparing treatment with palbociclib (a small molecule CDK4/6 inhibitor (CDK4/6i)) and interferon-gamma (IFN-γ), both known modulators of antigen presentation.^26,27^ Peptides derived from proteins implicated in the biological response to palbociclib and IFN-g were selectively enriched in the pMHC repertoire following treatment, connecting the intracellular response to extracellular immune presentation. Furthermore, peptides derived from the metabolic response to CDK4/6i, along with known tumor-associated antigens (TAAs), displayed significantly increased presentation with CDK4/6 inhibition. We propose this platform can be broadly applied to profile immunopeptidomic changes in a high-throughput, low-input format across sample types and treatments to inform combination therapy strategies and can be used to identify and quantify treatment-modulated antigen targets for targeted immunotherapy.

## RESULTS

### Quantitative immunopeptidomics platform using heavy isotope-coded MHC monomers improves quantitative accuracy

We set out to develop a platform to provide accurate relative and absolute quantification of pMHCs across multiple samples while controlling for losses associated with sample processing and enrichment. Accurate quantitative analysis is best performed with internal standards and multi-point internal calibration curves. To generate internal standards, heavy isotope labeled MHC peptides of interest were synthesized and loaded onto biotinylated MHC monomers through UV-mediated peptide exchange (Fig. 1a). To control for loading efficiency of synthetic peptides into recombinant MHC proteins, the concentration of stable hipMHC complexes was determined by an enzyme-linked immunosorbent assay (ELISA). Stable hipMHC complexes were then used in two ways: selected hipMHC complexes were spiked at the same concentration into whole cell lysate from each sample to provide a normalization correction for relative quantification across samples, while other hipMHC complexes were titrated at different concentrations into each sample to verify correction parameters, estimate dynamic range suppression for quantification, and/or create an internal standard curve for absolute quantification of a specific peptide. After adding hipMHCs, endogenous and exogenous pMHCs were isolated by immunoprecipitation, acid elution, and molecular weight size exclusion filtration.

**Figure 1.**
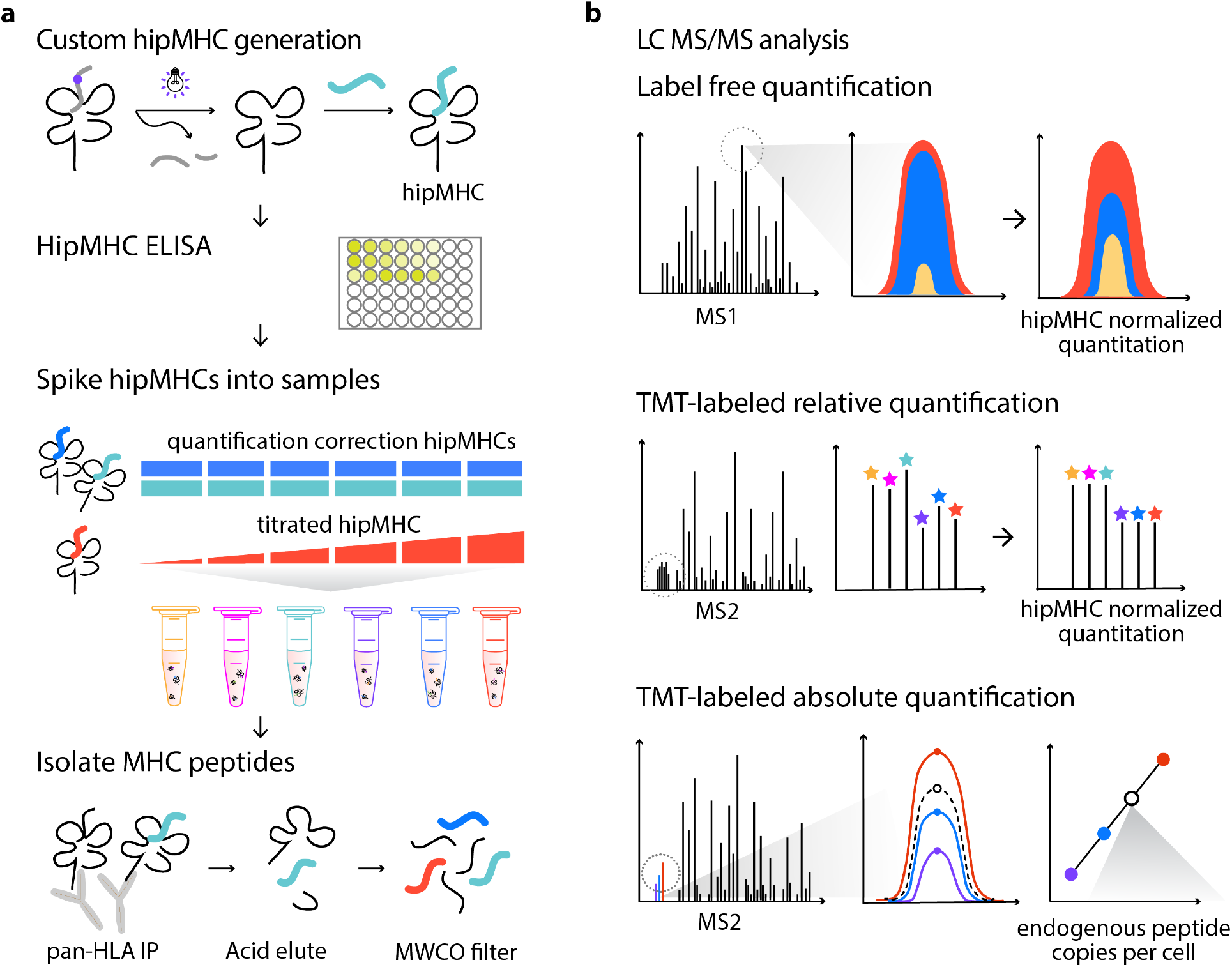
Platform for quantitative immunopeptidomics using hipMHCs for relative and absolute quantification. **a.** hipMHCs were generated through UV-mediated peptide exchange of HLA*A2:01 monomers with a heavy leucine HLA-A*02:01 binding peptide. Stable hipMHCs concentrations were measured with an ELISA, and hipMHC complexes were added to lysate samples prior to IP, at the same concentration for quantification correction (blue/teal), or titrated in to create an internal standard curve (red). Heavy and light pMHCs were isolated with immunoprecipitation, acid elution, and molecular weight cut-off (MWCO) filters. **b.** Peptides were analyzed by LC MS/MS three ways. Relative quantification label-free analyses were quantified by integrating the area under the curve (AUC) of the chromatographic elution across samples, and quantification was normalized by applying correction factors determined by hipMHC AUC intensity ratios between samples. Samples for multiplexed analysis were TMT-labeled and relative quantification was implemented using reporter ion intensities. Normalization was performed using hipMHC reporter ion intensity ratios across TMT channels. For absolute quantification, TMT-labeled samples containing a hipMHC internal standard curve were used to calculate the endogenous copies per cell of the pMHC of interest.

Peptide mixtures were next analyzed by liquid chromatography-tandem mass spectrometry (LC-MS/MS) in three different ways (Fig. 1b). For label-free (LF) analyses, samples were analyzed individually, and peptides were quantified by integrating the area under the curve (AUC) for the chromatographic elution of precursor masses for each peptide-spectrum match (PSM). Relative AUC intensities of quantification correction hipMHCs were used to normalize AUC intensities of endogenous peptides across analyses. To analyze multiple samples simultaneously, we labeled samples with tandem mass tags (TMT) and relative TMT ion intensity ratios of hipMHCs were used for normalization to correct the relative quantification in multiplexed samples. TMT-labeled titrated hipMHCs were also used for absolute quantification of endogenous peptides. Apex TMT intensities of hipMHCs generated a peptide specific multipoint calibration curve to calculate the average number of copies per cell. As a control, heavy isotope-coded synthetic peptides not complexed to MHC were spiked into whole cell lysate prior to immunoprecipitation. These peptides were not detected in the subsequent LC-MS/MS analysis, demonstrating that only peptides in stable complexes were isolated in our workflow and that excess free peptides did not displace endogenously-presented peptides.

### HipMHCs improve quantitative accuracy in label-free and tandem mass tag labeled samples

To demonstrate the improved quantitative accuracy obtained with hipMHCs, we used five LF and six TMT-labeled technical replicates of 1×10^7^ MDA-MB-231 breast cancer cells to measure variance between replicates before and after hipMHC correction (Fig. 2a). In both LF and TMT-labeled workflows, we spiked 30 fmol of two quantification correction hipMHCs into each sample, along with 30-300 fmol of a titrated hipMHC (Fig. 2b). A total of 2,369 unique pMHCs were identified in total across five LF analyses, 1,352 of which were quantifiable via AUC integration (Fig. 2c). Of these quantifiable peptides, only 589 were quantified in all five analyses, highlighting the poor run-to-run overlap of LF analyses, even with replicate samples (Supplementary Fig. 1a). By comparison, 1,754 unique peptides were quantifiable with TMT-labeled analyses. The extra sample handling steps associated with TMT labeling can result in losses, so to achieve high coverage of the immunopeptidome, labeled samples were divided into six separate analyses, thereby increasing the number of unique identifications (Supplementary Fig. 1b). In both LF and TMT analyses, peptides matched expected length distributions of 8 to 11 amino acids (Supplementary Fig. 1c), and 82% of LF and 92% of TMT 9-mers were predicted to be binders with <500nM predicted affinity (Fig. 2d).^28^ Further reducing the input material to 5×10^6^ cells still resulted in 86% of the number of unique peptides identified with 1×10^7^ cells, establishing the sensitivity of this method for low-input pMHC analyses. (Supplementary Fig. 1d).

**Figure 2.**
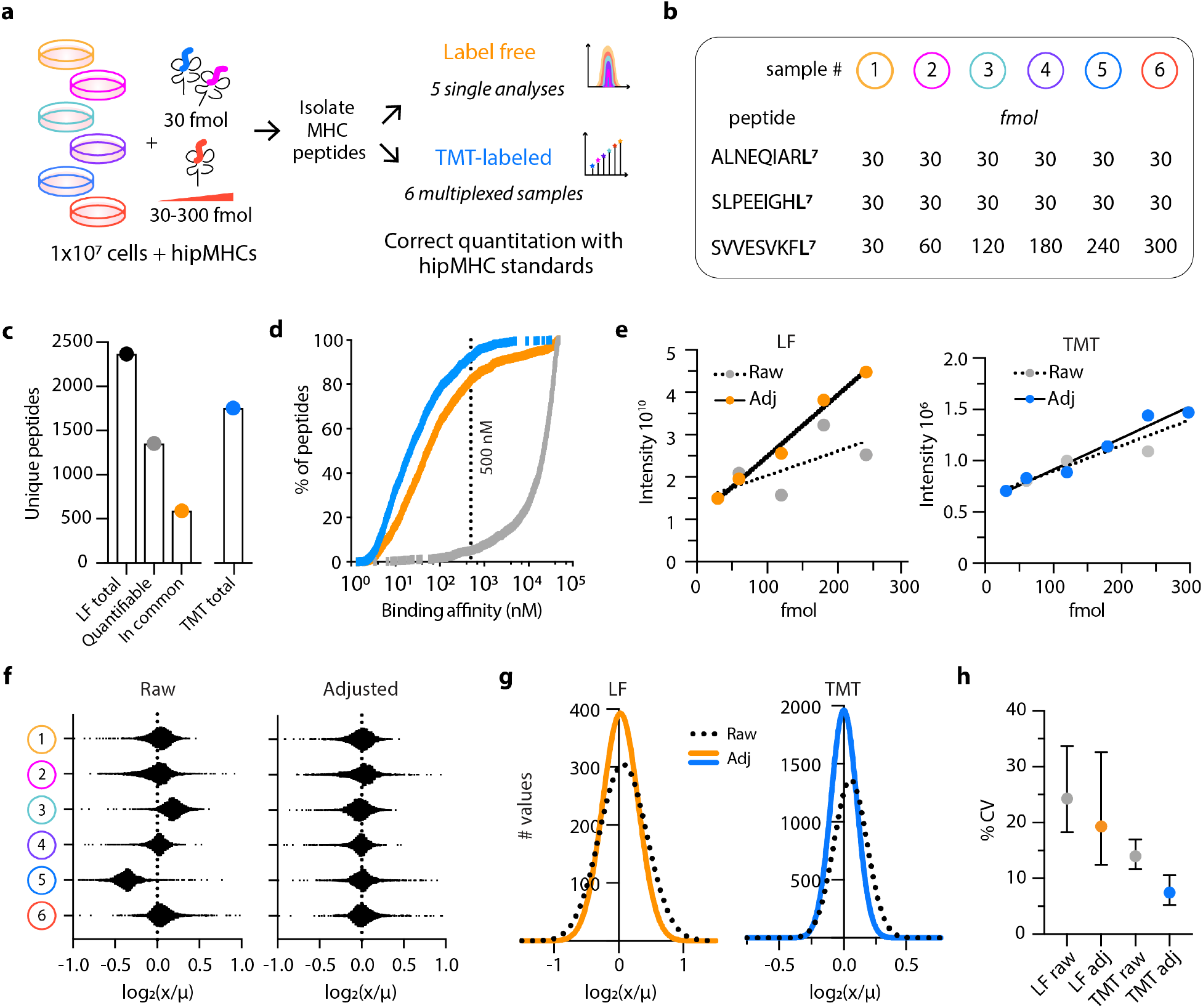
hipMHCs improve quantitation in LF and TMT-labeled samples. **a.** Experimental design. Five LF (orange) and six TMT (blue) technical replicates of 1×10^7^ cells + hipMHCs were used to compare LF & TMT quantification. **b.** Sequence and amount of hipMHC containing a heavy leucine (+7) peptide spiked into each sample. ALNEQIARL and SLEEPIGHL were used as quantification correction hipMHCs, and SVVESVKFL was titrated in across samples. For LF analysis, sample #6 was omitted. **c.** 2,369 unique LF peptides identified across five analyses (black), 1,352 of which were quantifiable (grey) via AUC quantification, and 589 quantifiable peptides which were identified in all five analyses (orange). 1,754 unique peptides were quantified with labeled analyses combining six TMT fractions (blue). **d.** 92% of TMT, 82% of LF, and 5% of random 9mers from the proteome (grey) are predicted to bind to an HLA allele in MDA-MB-231 cells with an affinity <500 nM. **e.** Linear fit of titrated hipMHC peptide for LF (left) and TMT (right). Raw r^2^ = 0.48 (LF) and r^2^=0.91 (TMT), hipMHC adjusted (adj) r^2^=0.99 (LF) and r^2^=0.96 (TMT). **f.** Distribution of the log_2_ fold change (FC) of each peptide’s quantification (x) over the mean (*µ*) peptide quantification across samples for raw (left) and hipMHC adjusted (right). **g.** Gaussian fit of the frequency distribution of log_2_ FC of (x) over (*µ*) for raw and hipMHC adjusted LF and TMT samples. 99.7% of variance between peptide quantitation (3x std dev) is captured within a 2.14 (raw) and 1.77 (adj) FC from the mean for LF samples, and 1.30 (raw) and 1.23 (adj.) for TMT samples. **h.** Median coefficient of variation (CV) for LF (24.29% raw, 19.32% adj) & TMT (14.00% raw, 7.48% adj). Error bars represent the interquartile range.

To normalize LF and TMT-labeled datasets, we applied correction parameters calculated from the quantification correction hipMHCs. The titrated peptide, SVVESVKFL^7^, displayed an improved linear fit after correction, with an even more pronounced effect in the LF samples (Fig. 2e). We observed dynamic range suppression for this peptide in TMT-labeled (4.7x) and LF (2.7x) datasets, demonstrating in both cases that quantitative differences are likely larger than what is measured.

In both analyses, hipMHC quantification correction reduced variation, for example, peptides from TMT-labeled sample 5 have lower intensities than the other samples, which was corrected by hipMHC normalization (Fig. 2f). The standard deviation from the mean for replicate samples decreased in LF and TMT-labeled samples (Fig. 2g), though TMT labeling showed lower variation between replicates (Fig. 2h), allowing for higher confidence in small shifts within the immunopeptidome.

### A hipMHC internal multipoint calibration curve enables absolute quantification of endogenous pMHCs

To demonstrate the ability of hipMHCs to quantify pMHC copies per cell, we selected two peptides identified in TMT-labeled MDA-MB-231 cells for absolute quantification: KLDVGNAEV derived from B cell receptor associated protein 31 (BCAP31) and KQVSDLISV from DEAD-box RNA helicase 5 (DDX5). BCAP31 regulates the transport of membrane proteins from the endoplasmic reticulum to the Golgi, a central component of antigen processing, and is a known TAA peptide.^29^ DDX5 is important in gene expression regulation and has been implicated in proliferation, metastasis, and tumorigenesis in cancer.^30^ These peptides were detected at differing levels with the highly abundant BCAP31 peptide falling in the 98^th^ percentile and DDX5 falling in the 33^rd^ percentile of abundance (Fig. 3a). Both peptides were synthesized with heavy-isotope labeled leucine (+7), and hipMHC normalization standards were added to three replicates of 1×10^7^ MDA-MB-231 cells along with titrated amounts of BCAP31 and DDX5 hipMHC (Fig. 3b). We then labeled samples with TMT and performed LC-MS/MS analysis using an inclusion list, so only targeted peptides of interest were selected for fragmentation.

**Figure 3.**
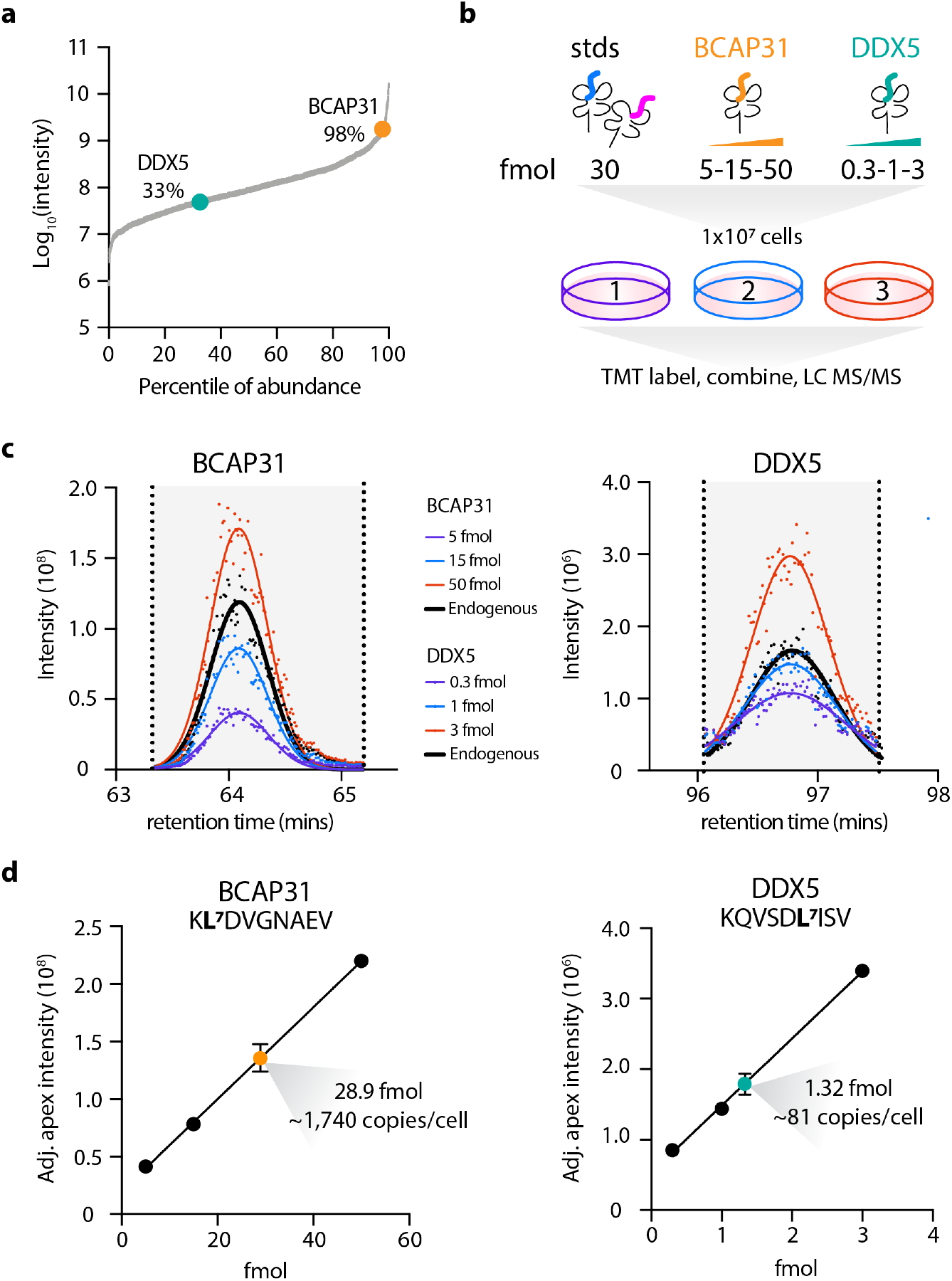
Absolute quantification of pMHCs using an internal hipMHC standard curve and isobaric labeling. **a.** Log_10_ value of average AUC intensity for TMT-labeled MDA-MB-231 cells from Fig. 2 determined by AUC quantification. Percentile of abundance represents a peptide’s rank relative to the most abundant peptide. **b.** Experimental design. Normalization standards along with 5, 15, and 50 fmol of BCAP31 and 0.3, 1, and 3 fmol of DDX5 hipMHCs were added to three replicates of 1×10^7^ MDA-MB-231 cells, and peptides from MHC complexes were isolated, labeled, and analyzed via LC-MS/MS. **c.** Chromatographic elution profiles for the three TMT reporter ion intensities of the hipMHC standard curve (colored), along with the average (n=3) TMT reporter ion intensity trace of the endogenous peptide (black). Each MS2 scan is represented as a single point, and elution profiles are fitted with a gaussian distribution (line). **d.** Adjusted (hipMHC normalized) apex intensity versus fmol of hipMHC added creates a standard curve from which the endogenous concentration of antigen is calculated. For both linear fits of BCAP31 and DDX5, r^2^ > 0.999. Error bars represent standard deviation between the 3 labeled replicates of the endogenous peptide.

Chromatographic traces of the three TMT reporter ions for heavy BCAP31 and DDX5 peptides displayed increasing ion intensities with increasing amount of hipMHC added (Fig. 3c). In order to quantify peptide expression, the apex intensities of reporter ions were adjusted based on the normalization hipMHCs, and a linear fit was used to pMHC concentration present in the sample. Cells had an average of 1,740 copies per cell of the BCAP31 peptide, and 81 copies per cell of the DDX5 peptide (Fig. 3d). Concentrations of the DDX5 hipMHC as low as 100 attomole were detected (six copies per cell) showcasing the broad range of pMHC expression levels quantifiable by our method (Supplementary Fig. 2). Furthermore, BCAP31 and DDX5 had a dynamic range suppression of 1.9x and 2.5x, respectively, illustrating that the ion suppression is not uniform across peptides and that peptide-specific internal standards may be required for absolute quantification of each pMHC of interest.

### CDK4/6 inhibition alters the pMHC repertoire in BRAF/NRAS mutant melanoma cells

Cyclin-dependent kinases 4 and 6 (CDK4/6) control cell cycle progression by phosphorylating Rb1, thereby releasing the E2F family of transcription factors that drive progression through the G1 checkpoint.^31^ CDK4/6 is often dysregulated and overactive in cancer, leading to uncontrolled proliferation.^32^ As such, CDK4/6 inhibitors have emerged as a potentially powerful class of anticancer agents. In recent years, CDK4/6i have also been shown to enhance tumor immunogenicity by increasing surface MHC class I expression and boosting T cell activation and infiltration.^27,33,34^ However, to date, the effect of CDK4/6 inhibition on the MHC class I peptide repertoire has not been characterized. We therefore applied our platform to quantify how pMHC repertoires change *in vitro* upon treatment with the CDK4/6i, palbociclib.

We selected four melanoma cell lines for analysis: SKMEL5 and SKMEL28 (BRAF mutant), and SKMEL2 and IPC298 (NRAS mutant). Based on sensitivity analyses for each cell line (Fig. 4a), we selected two doses of palbociclib for further study: a low dose of 1 μM, below the IC_50_ of all four cell lines, and a high dose of 10 μM, near the IC_50_. Three biological replicates of 1×10^7^ cells of each cell line were then treated with DMSO, low-dose, or high-dose CDK4/6i for 72h (Fig. 4b). Low-dose treatment increased surface class I MHC presentation, as measured by flow cytometry, by 1.5-2x across cell lines, whereas high-dose treatment had a milder effect (Fig. 4c, Supplementary Fig. 3a).

**Figure 4.**
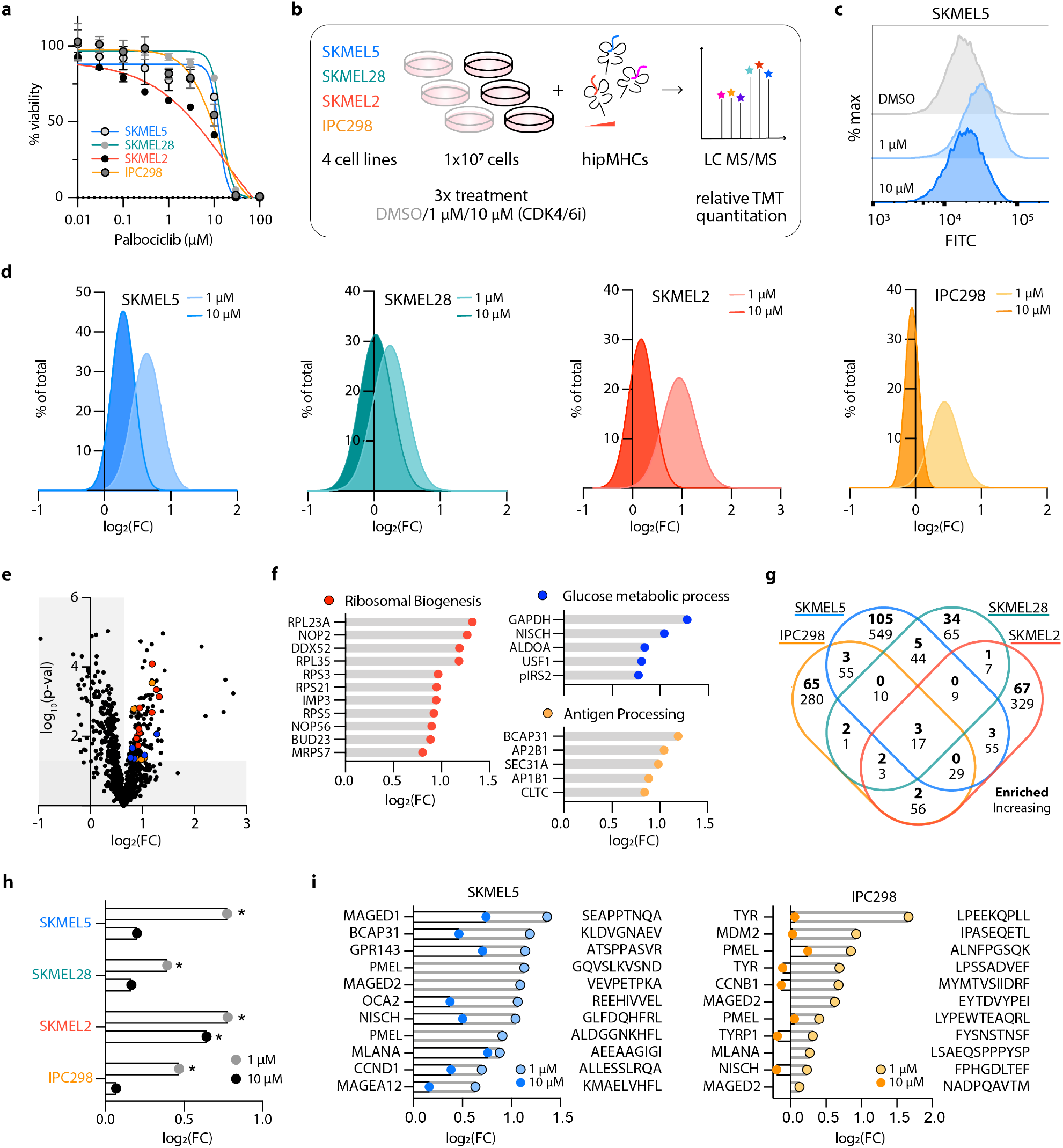
Quantitative immunopeptidomics reveals pMHC repertoire alterations with CDK4/6i in melanoma. **a.** Viability was assayed at 72h after drug treatment, data is represented as a fraction of the DMSO control. Calculated IC50s: for SKMEL5= 12.74 μM, SKMEL28=14.62 μM, SKMEL2=16.98 μM, IPC298=10.62 μM. **b.** Experimental setup. Three replicates of 1×10^7^ cells were treated with DMSO or CDK4/6i for 72h. hipMHC standards were added prior to immunoprecipitation for quantification correction, and isolated peptides were labeled with TMT and analyzed by MS. **c.** Flow cytometry measurements of surface HLA expression in SKMEL5 cells. Data is represented as % of maximum signal, and the distributions are representative of three replicates experiments. **d.** Histogram distribution of log_2_(fold change) (FC) of (CDK4/6i / DMSO) for unique pMHCs. Data is represented as a % of total unique peptides identified. **e.** Volcano plot displaying log_2_(FC) on x-axis, and significance on y-axis. Significance (paired t-test) was calculated after mean centering 1 μM treated data. Colored points correspond to the processes in **4f**. **f.** Log_2_(FC) of enriched peptides from GO term enrichment labeled with source protein. FDR q-value < 0.05. **g.** 4-way Venn-diagram of the number of source proteins of peptides significantly enriched significantly increasing. **h.** Log_2_(FC) of pIRS2 peptide following 1 μM (grey) or 10 μM (black) palbociclib, *p < 0.05. **i.** Gene, peptide sequence, and log_2_(FC) of TAAs in SKMEL5 (left) and IPC298 (right) cells.

To characterize the pMHC repertoire alterations induced by CDK4/6i, multiplexed relative quantitation was performed comparing low- and high-dose CDK4/6i to DMSO for each cell line, and data was normalized using hipMHC standards (Supplementary Fig. 3b). As with our previous analysis, identified peptides matched expected length distributions, and a majority were predicted to be MHC class I binders (Supplementary Fig. 3c-d). Immunopeptidomic analysis for each cell line and treatment showed a similar trend to the flow cytometry data: low-dose palbociclib shifted mean pMHC expression higher than DMSO treatment in all cell lines, and a high dose showed a small increase in mean expression for SKMEL5 compared to DMSO and no significant change for the other cell lines (Fig. 4e, Supplementary Fig. 3.e). We measured a wider distribution of changes in peptide presentation following low dose treatment, with several peptides increasing eight to ten-fold, even before considering the effect of dynamic range suppression (Supplementary Fig. 3b).

To gain insight into the biology underlying CDK4/6i modulated pMHC alterations, we analyzed our data in two ways. First, we determined which peptides and source proteins were significantly increased with CDK4/6 inhibition over DMSO. Because many peptides are significantly increased with low-dose treatment, we also identified the peptides and source proteins that were significantly enriched in presentation with treatment relative to the mean fold change of all peptides, highlighting peptides preferentially modulated by palbociclib.

Using these data, we performed GO term enrichment^35–37^ on peptides significantly enriched in low-dose treated SKMEL5 cells, and identified enriched biological processes of interest including ribosomal biogenesis, glucose metabolic process, and antigen processing, a reflection of the expected biological response to CDK4/6 inhibition (Fig. 4f).^27,38,39^ To determine if the measured pMHC alterations to SKMEL5 cells were common across cell lines, we compared the source proteins of peptides that were significantly enriched with low-dose treatment compared to DMSO across all four cell lines. Surprisingly, a majority (72-88%) of enriched source proteins were unique to each cell line, and we discovered only three proteins in common: vimentin (VIM), putative beta-actin-like protein 3 (POTEKP), and SIL1 nucleotide exchange factor (SIL1) (Fig. 3g, Supplementary Fig. 3f). Even when comparing source proteins of all peptides significantly increasing to any extent, just 17 proteins in common were identified, further illustrating the uniqueness of each immunopeptidomic landscape (Supplementary Fig. 3g). We investigated whether the commonality of these 17 proteins could be explained by having high abundance in the peptide mixtures, but in SKMEL5 cells they were scattered throughout the distribution of AUC intensities (Supplementary Fig. 3h).

While the list of shared pMHCs and source proteins in common is limited, of interest is the serine-phosphorylated IRS2 (pIRS2) peptide, RVApSPTSGVK. This post-translationally modified sequence has previously been shown to be restricted to malignant cells, with only the phosphorylated form demonstrating immunogenic potential^40,41^ Despite differences in the cell lines’ allelic profiles, was observed the pIRS2 peptide increasing across all cell lines with low dose treatment (Fig. 3h). Furthermore, RVApSPTSGVK has high expression among pMHCs (Supplementary Fig 3h), and can be isolated without phospho-enrichment.^42^ As a result, this peptide may be uniquely positioned as a broadly targetable antigen whose expression can be modulated by CDK4/6 inhibition. As a general effect of CDK4/6i treatment, TAAs derived from proteins like MLANA (MART1), PMEL (gp100), and TYR, among others, also increased in presentation following treatment (Fig. 4i). While these antigens and their source proteins are not universally conserved across our cell lines, the effect of increased TAA presentation following 1μM palbociclib treatment could be applied to increase antigen presentation prior to immunotherapies targeting these well-documented antigens.

### Cellular response to CDK4/6i is reflected in quantitative differences in immunopeptidome

To further assess whether quantitative differences in the immunopeptidome after CDK4/6 inhibition are reflective of the cell signaling response to a perturbation, we performed a nonparametric test to identify positively- and negatively-enriched pathways. Gene names for source proteins were rank ordered according to fold change with treatment and searched against the MSigDB Hallmarks gene set database using Gene Set Enrichment Analysis (GSEA).^43–45^ This analysis did not reveal any significantly enriched pathways for the low dose treatment, but high dose CDK4/6 inhibition showed significant enrichment among downregulated pMHCs of E2F targets, G2M checkpoint, DNA repair, mitotic spindle, and MTORC1 signaling pathways in one or more cell lines (Fig. 5a). These findings reflect the known biological effects of CDK4/6i. For instance, inhibiting CDK4/6 decreases expression of E2F targets, and peptides derived from E2F targets like Ki-67, a proliferation marker, were depleted in all four cell lines (Fig. 5b). E2F also controls genes involved in DNA damage repair, and consistently, γH2AX levels, a marker of DNA double-strand breaks, increased at 72h with CDK4/6i treatment in a dose dependent manner (Supplemental Fig. 4a).^46^ Although similar biological processes are enriched across the four cell lines, source proteins for significantly depleted E2F peptides showed little overlap between the cell lines (Fig. 5c), again emphasizing the individuality of each cell line’s detected pMHC repertoire.

**Figure 5.**
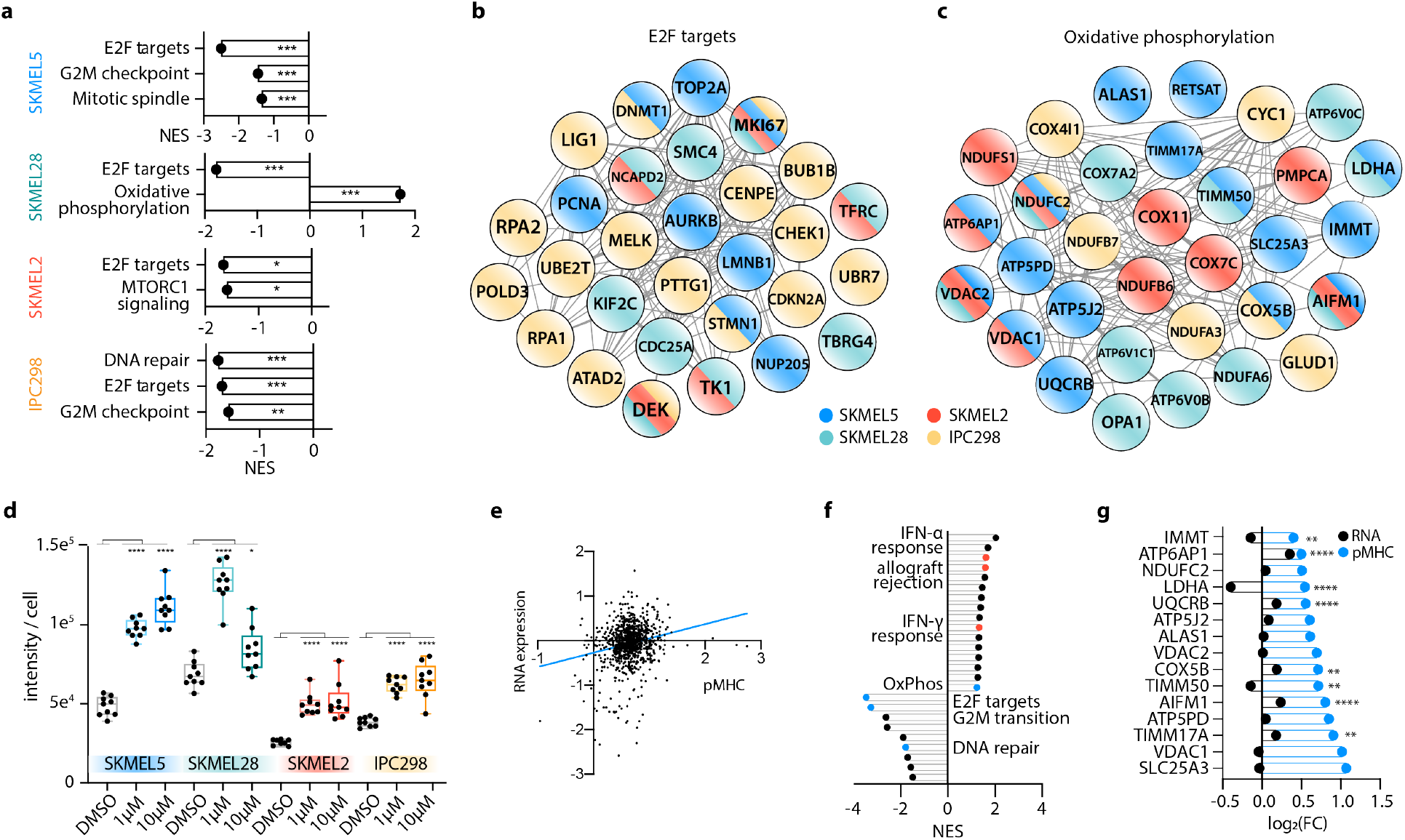
Pathway analysis of CDK4/6i-altered immunopeptidome aligns with the cellular response to treatment. **a.** Enrichment analysis of high dose CDK4/6i treated cell lines. Significantly enriched pathways (q < 0.25, p < 0.05) are reported, and y-axis indicates the normalized enrichment score (NES) where +/− scores reflect enrichment directionality. *p < 0.05, **p < 0.01, ***p < 0.001. **b-c.** String network of protein-protein interactions of all source proteins from E2F peptides **(b)** significantly decreasing with 10 μM CDK4/6i, and OxPhos peptides **(c)** significantly increasing with 1 μM CDK4/6i for all cell lines except SKMEL28, where peptides from 10 μM are depicted. Node color corresponds to cell line. **d.** Quantification (n=9) of MitoTraker green intensity normalized to cell number following 72h CDK4/6i treatment. Data is represented as a box and whiskers plot with whiskers displaying minimum and maximum signal. Significance was determined using a one-way ANOVA for each condition vs. DMSO. *p < 0.05, ****p < 0.0001. **e.** The relationship between log_2_(FC) of CDK4/6i / DMSO for RNA expression (y-axis) and pMHC presentation (x-axis) of SKMEL5 cells treated for 72h with 1 μM CDK4/6i, r^2^ = 0.04. **f**. GSEA against hallmark gene sets using RNA-seq data. Annotated pathways reflect pathways also identified in the immunopeptidome analysis (blue), and those that match with previous reported data (red).^1^ **g.** Log_2_(FC) for SKMEL5 OxPhos peptides significantly increasing (blue) with CDK4/6i, and matched RNA expression (black). P-values indicate pMHC significance (CDK4/6i vs. DMSO), **p < 0.01, ****p < 0.0001.

Only one pathway, oxidative phosphorylation (OxPhos), was significantly upregulated in SKMEL28 cells. However, all cell lines presented peptides derived from the OxPhos pathway that increased significantly with CDK4/6 inhibition, especially at low-dose treatment (Fig. 5c). OxPhos has been shown to increase with CDK4/6i due to increased ATP levels and mitochondrial mass, elevating metabolic activity. Comparably, all samples showed elevated mitochondrial levels following treatment, suggesting that enriched pMHC presentation of OxPhos derived peptides reflects a change in the metabolic cell state (Fig. 5d).

Because alterations to the pMHC repertoire align with previously characterized biological responses to CDK4/6 inhibition, we tested whether changes in RNA expression could predict the quantitative immunopeptidome changes. No bulk correlation (r^2^=0.04) was observed between pMHC expression and RNA expression (Fig. 5e). This was unsurprising, as many mechanisms beyond gene expression regulate pMHC presentation, including protein synthesis, degradation, post-transitional modifications, processing, and more. Despite this poor correlation, significantly enriched gene sets in the immunopeptidome were also present in our RNA sequencing analysis (Fig. 5f). While E2F pMHCs significantly depleted in SKMEL5 cells correlated with significantly decreased gene expression of the same source proteins, (Supplementary fig. 4b), only five of the 15 positively enriched OxPhos peptides displayed significantly higher gene expression with CDK4/6 inhibition, with three decreasing in expression, and seven remaining unchanged (Fig. 5g). Collectively, these data suggest that while changes in gene expression and pMHC repertoires map to the same biological pathways, individual gene expression changes are not necessarily predictive of alterations in the immunopeptidome.

### Repertoire changes induced by IFN-γ stimulation are distinct from those induced by CDK4/6 inhibition

Previous work has demonstrated that CDK4/6 inhibition stimulates interferon signaling, augmenting antigen presentation levels.^15^ We also observed upregulation of IFN-γ response genes with low-dose CDK4/6i, as well as increased expression of genes relating to antigen presentation (Fig. 5f, Fig. 6a). Consequently, we tested whether direct IFN-γ stimulation would shift the repertoire similarly to CDK4/6 inhibition. Cells were stimulated with DMSO or 10 ng/mL IFN-γ for 72h and the resulting pMHC repertoires were quantified using our multiplexed hipMHC platform. IFN-γ increased surface pMHC levels greater than 2x for each cell line, a trend that was reflected in the immunopeptidome, as nearly every identified pMHC increased in presentation with stimulation (Fig. 6b, Supplementary Fig. 5a).

**Figure 6.**
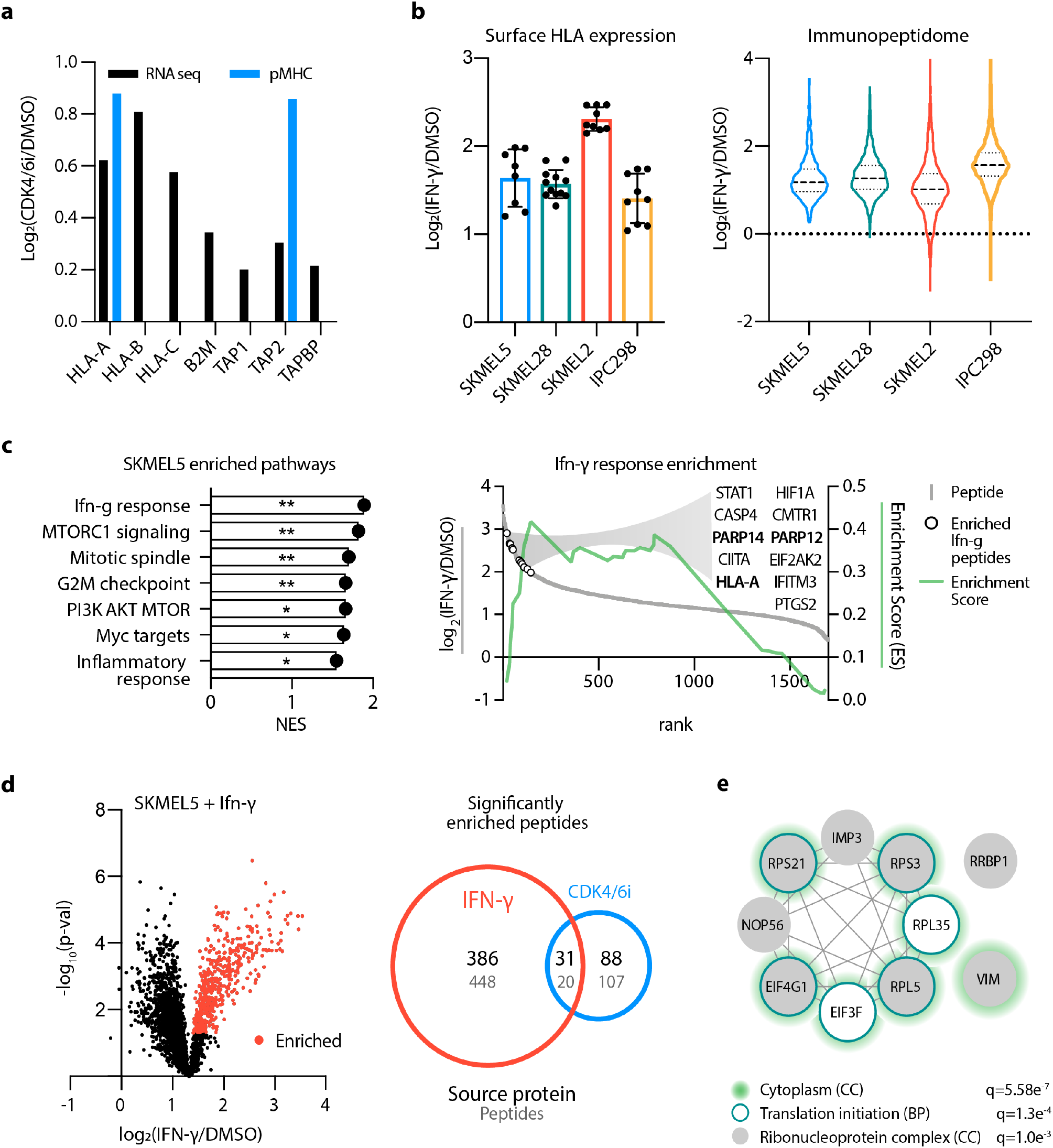
Quantifying the pMHC repertoire response to IFN-γ stimulation. **a**. Enrichment of antigen processing genes with 1μM CDK4/6 inhibition. RNA-seq (black) and pMHC (blue) values are shown as the log_2_ fold change (FC) of (IFN-γ/ DMSO). **b.** Surface HLA expression via flow cytometry (left) of cells treated with 10ng/mL IFN-γ for 72h shown as log_2_(FC). Errors bars represent standard deviation (n=8-12). Immunopeptidome log2(FC) (right), Dashed lines display quartiles, and mean fold changes are 2.42, 2.50, 2.08, and 3.04 for SKMEL5, SKMEL28, SKMEL2, and IPC298 cells, respectively. **c.** Enrichment analysis (left) of SKMEL5 cells treated with 10ng/mL IFN-γ for 72h. Y-axis indicates normalized enrichment score, q < 0.25, *p < 0.05, **p < 0.01. Enrichment plot (right) displays running enrichment score (green, right y-axis), and the log_2_(FC) (left y-axis) vs. rank (x-axis) for each ranked peptide (grey). Open circles show significantly enriched IFN-γ peptides. **d**. Volcano plot (left) of IFN-γ induced changes in SKMEL5 cells. Peptides are presented as the log_2_(FC) versus mean adjusted p-value. Red points represent peptides significantly enriched, (p < 0.05, fold change > 2.42). Venn diagram (right) of significantly enriched source proteins (black) or peptides (grey) between IFN-γ and 1μM CDK4/6i treated SKMEL5 cells. **e.** Protein-protein interaction network of source proteins significantly enriched in common. Gene ontology analysis for cellular components (CC) and biological professes (BP) annotate nodes as components of the cytoplasm and ribonucleoprotein complexes, or translation initiation. Significance values are FDR-corrected.

To determine the similarity of response to CDK4/6i, we again performed GSEA against the hallmark gene sets. The most significantly upregulated pathway in SKMEL5 cells with IFN-γ stimulation was the “IFN-γ response” (Fig. 6c). In fact, IFN-γ response was the top enriched pathway in every cell line, reiterating that the cellular response to stimulus is reflected in quantitative differences in pMHC presentation, and that IFN-γ related peptides are preferentially upregulated by IFN-γ stimulation (Supplementary Fig. 5b). Other pathways such as G2M checkpoint and mitotic spindle were positively enriched in IFN-γ stimulated cells, in contrast to the results of CDK4/6 inhibition.

Although the cell lines showed differential pMHC pathway enrichment upon CDK4/6i and IFN-γ stimulation, we tested whether any pMHCs or source proteins were commonly enriched in response to CDK4/6i and IFN-γ. In SKMEL5 cells we identified just 20 peptides and 31 source proteins significantly enriched in both conditions (Fig. 6d), which primarily map to the cytoplasm and contain multiple ribosomal and translation initiation proteins frequently overrepresented in immunopeptidomic datasets (i.e., DRiPs) (Fig. 6e).^47^ These data demonstrate that while CDK4/6 inhibition may induce an IFN-γ response, stimulating cells with IFN-γ does not recapitulate the distinct peptide repertoire alterations observed with CDK4/6i. Instead, IFN-γ stimulation alters the repertoire by augmenting the presentation of IFN-γ related peptides.

## Discussion

The addition of hipMHCs as internal standards improves relative quantitative accuracy for both LF and multiplexed labeled analyses, though multiplexed labeling with TMT showed superior accuracy and peptide binding specificity and yielded a higher number of quantifiable unique peptides using equivalent sample input. These internal hipMHC standards, which travel through the entire pMHC workflow, also account for variation across samples and provide an estimate for dynamic range suppression, which varies across peptides. While we use TMT 6-plex in our analyses, this method is compatible with other isobaric labeling strategies, including iTRAQ, TMT 11-plex and TMTpro, to analyze up to 16 samples simultaneously. For rapid profiling of immunopeptidome changes, we elected to use minimal sample input, making this protocol easily translatable for in vivo-derived tissue (*e.g.*, clinical and animal) samples. However, using this same general platform of hipMHCs and isobaric multiplex labeling, sample amount could be increased and coupled with fractionation for deeper sequencing of the pMHC repertoire.^48^

In addition to improved relative quantification, we also demonstrated the utility of hipMHCs for pMHC absolute quantification by generating an embedded multipoint standard curve. Using targeted mass spectrometry to detect attomole levels of antigen from just 1×10^7^ cells and regressing this signal against the titrated hipMHC standard, we were able to extract accurate absolute quantification in terms of copies per cell for two pMHC’s with ~20-fold difference in abundance. The ability to readily determine the absolute quantification of any detectable antigen of interest without the need for a pMHC-specific antibody will aid in targeted immunotherapy design. For instance, peptides of lower abundances may be better suited for engineered TCR-based therapies, as TCRs have been shown to be incredibly sensitive with as few as one pMHC complex being capable of initiating detectable T cell activation.^49^ Alternatively, antibody-based therapies targeting specific pMHCs, *e.g.*, bi-specific T cell engagers (BiTES) or antibody-drug conjugates (ADCs), may benefit from higher antigen expression levels, though results vary across antigen targets and antibody affinities.^50^ Moreover, absolute quantification of pMHC expression can help to untangle the biological relationships among antigen processing, epitope abundances, immunogenicity, and off-target toxicity (*e.g.*, tumor vs. non-tumor abundance).

It is worth noting that one existing restriction to using hipMHCs is the commercial availability of UV-mediated MHC monomers and ELISA control reagents, which are limited to a handful of common human class I alleles. While matched allele hipMHCs are not required for normalization correction if MHC molecules are isolated using a pan-specific antibody, they are necessary for accurate absolute quantification with embedded standard curves. An analogous technology, disulfide-stabilized HLA molecules, could be used in place of UV-mediated exchange.^51^ These HLA-B2M complexes show increased stability and higher exchange efficiency of lower-affinity peptides, potentially eliminating the need for an ELISA to quantify exchange efficiency and simplifying MHC refolding to expand this protocol to other alleles and species.

We applied our quantitative multiplexed hipMHC normalization to determine the pMHC repertoire response to CDK4/6 inhibition in melanoma. These results indicate that extracellular changes in pMHC abundance are reflective of the intracellular response to CDK4/6i. Moreover, CDK4/6i increased the presentation of TAAs and peptides derived from metabolic processes. Recently, high tumor antigen and metabolic protein expression levels have been shown to be predictive of checkpoint inhibitor response in melanoma, suggesting that CDK4/6i could be used in conjunction with checkpoint blockade or TIL-based therapies to increase tumor immunogenicity.^52^ As an alternate therapeutic strategy, peptide antigens whose surface expression was selectively increased by CDK4/6i could be utilized for targeted immunotherapy, either alone or in combination.

Indeed, the landscape of clinical trials exploring combination treatment regimens coupling checkpoint blockade with other therapies is rapidly expanding.^53–57^ Quantifying the molecular consequences of these combination regimes with our platform could provide novel insights into these trials and enable the informed design of new therapeutic combinations, potentially with targeted immunotherapies. Taken together, our relative and absolute quantitative immunopeptidomic data demonstrate the utility of quantitative immunopeptidomics in evaluating the pMHC repertoire response to therapy. The multiplexed nature of this platform allows for analyses of many samples in a short timescale, an important feature in the context of clinical trials. Further analyses of pMHC repertoire changes will be useful in understanding the order and timing of therapies to achieve optimal success and may enable predictions as to how to tune the immunopeptidome to be most applicable to immunotherapy targeting.

## Acknowledgements

We thank Array Biopharma for providing melanoma cell lines; Cristine Devlin and Michael Birnbaum for providing recombinant MHC monomers; Joe Card (Marto lab, Dana-Farber) for kindly providing guidance on the SP3 protocol; the MIT BioMicro Center (Stuart Levine) and the Swanson Biotechnology Center for technical support, specifically the Flow Cytometry (Glenn Paradis), Biopolymers & Proteomics (Richard Cook & Antonius Koller), and Bioinformatics (Charlie Whittaker) core facilities. This research was supported by funding from NIH Grants U54 CA210180, P30 CA14051, Melanoma Research Alliance Team Science Award #565436, the MIT Center for Precision Cancer Medicine, The Koch Institute Frontier Research Program, the Kathy and Curt Marble Cancer Research Fund, and a generous gift from Takeda Pharmaceuticals Immune Oncology Research Fund. L.E. Stopfer is supported by an NIH Training Grant in Environmental Toxicology (# T32-ES007020).

## Contributions

LES and FMW conceived the project. FMW and DAL advised the project. LES and JMM performed experiments. LS and BAJ analyzed data. LS, BAJ, and FMW wrote the manuscript, and JMM and DAL provided manuscript feedback.

## REFERENCES

1. Sharma, P., Hu-Lieskovan, S., Wargo, J. A. & Ribas, A. Leading Edge Review Primary, Adaptive, and Acquired Resistance to Cancer Immunotherapy. doi:10.1016/j.cell.2017.01.017.

2. Martins, F. et al. Adverse effects of immune-checkpoint inhibitors: epidemiology, management and surveillance. Nat. Rev. Clin. Oncol. doi:10.1038/s41571-019-0218-0.

3. Zappasodi, R., Merghoub, T. & Wolchok, J. D. Cancer Cell Perspective Emerging Concepts for Immune Checkpoint Blockade-Based Combination Therapies. doi:10.1016/j.ccell.2018.03.005.

4. Reits, E. A. et al. Radiation modulates the peptide repertoire, enhances MHC class I expression, and induces successful antitumor immunotherapy. J. Exp. Med. (2006) doi:10.1084/jem.20052494.

5. Brea, E. J. et al. Kinase Regulation of Human MHC Class I Molecule Expression on Cancer Cells. Cancer Immunol Res 4, 936–47.

6. Liu, L. et al. The BRAF and MEK Inhibitors Dabrafenib and Trametinib: Effects on Immune Function and in Combination with Immunomodulatory Antibodies Targeting PD-1, PD-L1, and CTLA-4. Clin. Cancer Res. 21, (2015).

7. Khallouf, H. et al. 5-Fluorouracil and Interferon-α Immunochemotherapy Enhances Immunogenicity of Murine Pancreatic Cancer Through Upregulation of NKG2D Ligands and MHC Class I. J. Immunother. 35, 245–253 (2012).

8. Liu, W. M., Fowler, D. W., Smith, P. & Dalgleish, A. G. Pre-treatment with chemotherapy can enhance the antigenicity and immunogenicity of tumours by promoting adaptive immune responses. Br. J. Cancer 102, 115–123 (2010).

9. He, S. et al. Enhanced interaction between natural killer cells and lung cancer cells: Involvement in gefitinib-mediated immunoregulation. J. Transl. Med. 11, (2013).

10. Sullivan, R. J. et al. Atezolizumab plus cobimetinib and vemurafenib in BRAF-mutated melanoma patients. Nature Medicine vol. 25 929–935 (2019).

11. Ascierto, P. A. et al. Dabrafenib, trametinib and pembrolizumab or placebo in BRAF-mutant melanoma. Nature Medicine vol. 25 941–946 (2019).

12. Hunt, D. et al. Characterization of peptides bound to the class I MHC molecule HLA-A2.1 by mass spectrometry. Science (80-.). 255, (1992).

13. Bassani-Sternberg, M., Pletscher-Frankild, S., Jensen, L. J. & Mann, M. Mass spectrometry of human leukocyte antigen class I peptidomes reveals strong effects of protein abundance and turnover on antigen presentation. Mol. Cell. Proteomics 14, 658–73 (2015).

14. Khodadoust, M. S. et al. Antigen presentation profiling reveals recognition of lymphoma immunoglobulin neoantigens. Nature 543, 723–727 (2017).

15. Bogunovic, B., Srinivasan, P., Ueda, Y., Tomita, Y. & Maric, M. Comparative Quantitative Mass Spectrometry Analysis of MHC Class II-Associated Peptides Reveals a Role of GILT in Formation of Self-Peptide Repertoire. PLoS One 5, 10599 (2010).

16. Shetty, V. et al. Quantitative immunoproteomics analysis reveals novel MHC class I presented peptides in cisplatin-resistant ovarian cancer cells. J. Proteomics 75, 3270–3290 (2012).

17. Jensen, S. M., Potts, G. K., Ready, D. B. & Patterson, M. J. Specific MHC-I Peptides Are Induced Using PROTACs. Front. Immunol. 9, (2018).

18. Loffler, M. W. et al. Mapping the HLA ligandome of colorectal cancer reveals an imprint of malignant cell transformation. Cancer Res. 78, 4627–4641 (2018).

19. Murphy, J. P. et al. Multiplexed Relative Quantitation with Isobaric Tagging Mass Spectrometry Reveals Class I Major Histocompatibility Complex Ligand Dynamics in Response to Doxorubicin. Anal. Chem. 91, 5106–5115 (2019).

20. Schittenhelm, R. B., Sian, T. C. C. L. K., Wilmann, P. G., Dudek, N. L. & Purcell, A. W. Revisiting the Arthritogenic Peptide Theory: Quantitative Not Qualitative Changes in the Peptide Repertoire of HLA-B27 Allotypes. Arthritis Rheumatol. 67, 702–713 (2015).

21. Hassan, C. et al. Accurate quantitation of MHC-bound peptides by application of isotopically labeled peptide MHC complexes. J. Proteomics 109, 240–244 (2014).

22. Hogan, K. T. et al. Use of selected reaction monitoring mass spectrometry for the detection of specific MHC class I peptide antigens on A3 supertype family members. Cancer Immunol. Immunother. 54, 359–371 (2005).

23. Tan, C. T., Croft, N. P., Dudek, N. L., Williamson, N. A. & Purcell, A. W. Direct quantitation of MHC-bound peptide epitopes by selected reaction monitoring. Proteomics 11, 2336–2340 (2011).

24. Wu, T. et al. Quantification of epitope abundance reveals the effect of direct and cross-presentation on influenza CTL responses. doi:10.1038/s41467-019-10661-8.

25. Bozzacco, L. et al. Mass spectrometry analysis and quantitation of peptides presented on the MHC II molecules of mouse spleen dendritic cells. J. Proteome Res. 10, 5016–5030 (2011).

26. Yang, Y., Xiang, Z., Ertl, H. C. J. & Wilson, J. M. Upregulation of class I major histocompatibility complex antigens by interferon γ is necessary for T-cell-mediated elimination of recombinant adenovirus-infected hepatocytes in vivo. Proc. Natl. Acad. Sci. U. S. A. 92, 7257–7261 (1995).

27. goel, shom et al. CDK4/6 inhibition triggers anti-tumour immunity. Nat. Publ. Gr. 548, (2017).

28. Jurtz, V. et al. NetMHCpan-4.0: Improved Peptide–MHC Class I Interaction Predictions Integrating Eluted Ligand and Peptide Binding Affinity Data. J. Immunol. 199, 3360–3368 (2017).

29. Gloger, A., Ritz, D., Fugmann, T. & Neri, D. Mass spectrometric analysis of the HLA class I peptidome of melanoma cell lines as a promising tool for the identification of putative tumor-associated HLA epitopes Europe PMC Funders Group. Cancer Immunol Immunother 65, 1377–1393 (2016).

30. Nyamao, R. M., Wu, J., Yu, L., Xiao, X. & Zhang, F. M. Roles of DDX5 in the tumorigenesis, proliferation, differentiation, metastasis and pathway regulation of human malignancies. Biochimica et Biophysica Acta - Reviews on Cancer vol. 1871 85–98 (2019).

31. Choi, Y. J. & Anders, L. Signaling through cyclin D-dependent kinases. Oncogene vol. 33 1890–1903 (2014).

32. Hamilton, E. & Infante, J. R. Targeting CDK4/6 in patients with cancer. Cancer Treatment Reviews vol. 45 129–138 (2016).

33. Deng, J. et al. CDK4/6 Inhibition Augments Antitumor Immunity by Enhancing T-cell Activation. (2017) doi:10.1158/2159-8290.CD-17-0915.

34. Chaikovsky, A. C. & Sage, J. Beyond the cell cycle: Enhancing the immune surveillance of tumors via CDK4/6 inhibition. Molecular Cancer Research vol. 16 1454–1457 (2018).

35. Szklarczyk, D. et al. STRING v11: Protein-protein association networks with increased coverage, supporting functional discovery in genome-wide experimental datasets. Nucleic Acids Res. 47, D607–D613 (2019).

36. Ashburner, M. et al. Gene ontology: Tool for the unification of biology. Nature Genetics vol. 25 25–29 (2000).

37. The Gene Ontology Resource: 20 years and still GOing strong. Nucleic Acids Res. 47, D330–D338 (2019).

38. Donati, G., Montanaro, L. & Derenzini, M. Ribosome biogenesis and control of cell proliferation: p53 is not alone. Cancer Research vol. 72 1602–1607 (2012).

39. Franco, J., Balaji, U., Freinkman, E., Witkiewicz, A. K. & Knudsen, E. S. Metabolic re-programming of pancreatic cancer mediated by CDK4/6 inhibition elicits unique vulnerabilities. doi:10.1016/j.celrep.2015.12.094.

40. Zarling, A. L. et al. Identification of class I MHC-associated phosphopeptides as targets for cancer immunotherapy. Proc. Natl. Acad. Sci. U. S. A. 103, 14889–94 (2006).

41. Zarling, A. L. et al. MHC-restricted phosphopeptides from insulin receptor substrate-2 and CDC25b offer broad-based immunotherapeutic agents for cancer. Cancer Res. 74, 6784–95 (2014).

42. Bassani-Sternberg, M. et al. Direct identification of clinically relevant neoepitopes presented on native human melanoma tissue by mass spectrometry. Nat. Commun. 7, 13404 (2016).

43. Mootha, V. K. et al. PGC-1α-responsive genes involved in oxidative phosphorylation are coordinately downregulated in human diabetes. Nat. Genet. 34, 267–273 (2003).

44. Subramanian, A. et al. Gene set enrichment analysis: A knowledge-based approach for interpreting genome-wide expression profiles. Proc. Natl. Acad. Sci. U. S. A. 102, 15545–15550 (2005).

45. Liberzon, A. et al. The Molecular Signatures Database Hallmark Gene Set Collection. Cell Syst. 1, 417–425 (2015).

46. Hashizume, R. et al. Inhibition of DNA damage repair by the CDK4/6 inhibitor palbociclib delays irradiated intracranial atypical teratoid rhabdoid tumor and glioblastoma xenograft regrowth. Neuro. Oncol. 18, 1519–1528 (2016).

47. Bourdetsky, D., Schmelzer, C. E. H. & Admon, A. The nature and extent of contributions by defective ribosome products to the HLA peptidome. doi:10.1073/pnas.1321902111.

48. Purcell, A. W., Ramarathinam, S. H. & Ternette, N. Mass spectrometry–based identification of MHC-bound peptides for immunopeptidomics. Nat. Protoc. 14, 1687–1707 (2019).

49. Huang, J. et al. Article A Single Peptide-Major Histocompatibility Complex Ligand Triggers Digital Cytokine Secretion in CD4 + T Cells. Immunity 39, 846–857 (2013).

50. Ellerman, D. Bispecific T-cell engagers: Towards understanding variables influencing the in vitro potency and tumor selectivity and their modulation to enhance their efficacy and safety. Methods 154, 102–117 (2019).

51. Moritz, A. et al. High-throughput peptide-MHC complex generation and kinetic screenings of TCRs with peptide-receptive HLA-A*02:01 molecules. Sci. Immunol. 4, (2019).

52. Harel, M. et al. Proteomics of Melanoma Response to Immunotherapy Reveals Mitochondrial Dependence. doi:10.1016/j.cell.2019.08.012.

53. Esteva, F. J., Hubbard-Lucey, V. M., Tang, J. & Pusztai, L. Immunotherapy and targeted therapy combinations in metastatic breast cancer. The Lancet Oncology vol. 20 e175–e186 (2019).

54. Yu, C. et al. Combination of immunotherapy with targeted therapy: Theory and practice in metastatic melanoma. Frontiers in Immunology vol. 10 (2019).

55. McGranahan, T., Therkelsen, K. E., Ahmad, S. & Nagpal, S. Current State of Immunotherapy for Treatment of Glioblastoma. Current Treatment Options in Oncology vol. 20 (2019).

56. Ghisoni, E., Imbimbo, M., Zimmermann, S. & Valabrega, G. Ovarian cancer immunotherapy: Turning up the heat. Int. J. Mol. Sci. 20, (2019).

57. George, S., Rini, B. I. & Hammers, H. J. Emerging Role of Combination Immunotherapy in the First-line Treatment of Advanced Renal Cell Carcinoma: A Review. JAMA Oncology vol. 5 411–421 (2019).

