## Supplemental Figures for "Multiplexed relative and absolute quantitative immunopeptidomics reveals MHC I repertoire alterations induced by CDK4/6 inhibition"

### Supplemental figure 1

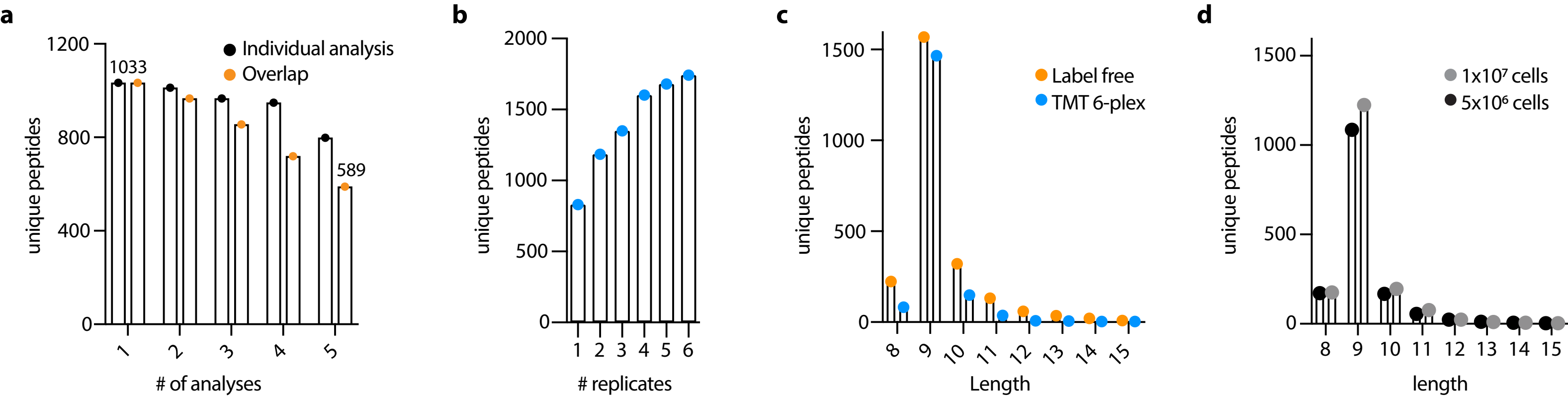

Supplemental figure 2

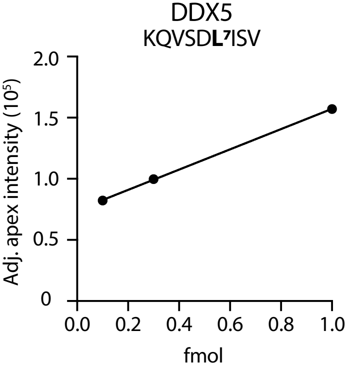

Supplementary figure 3

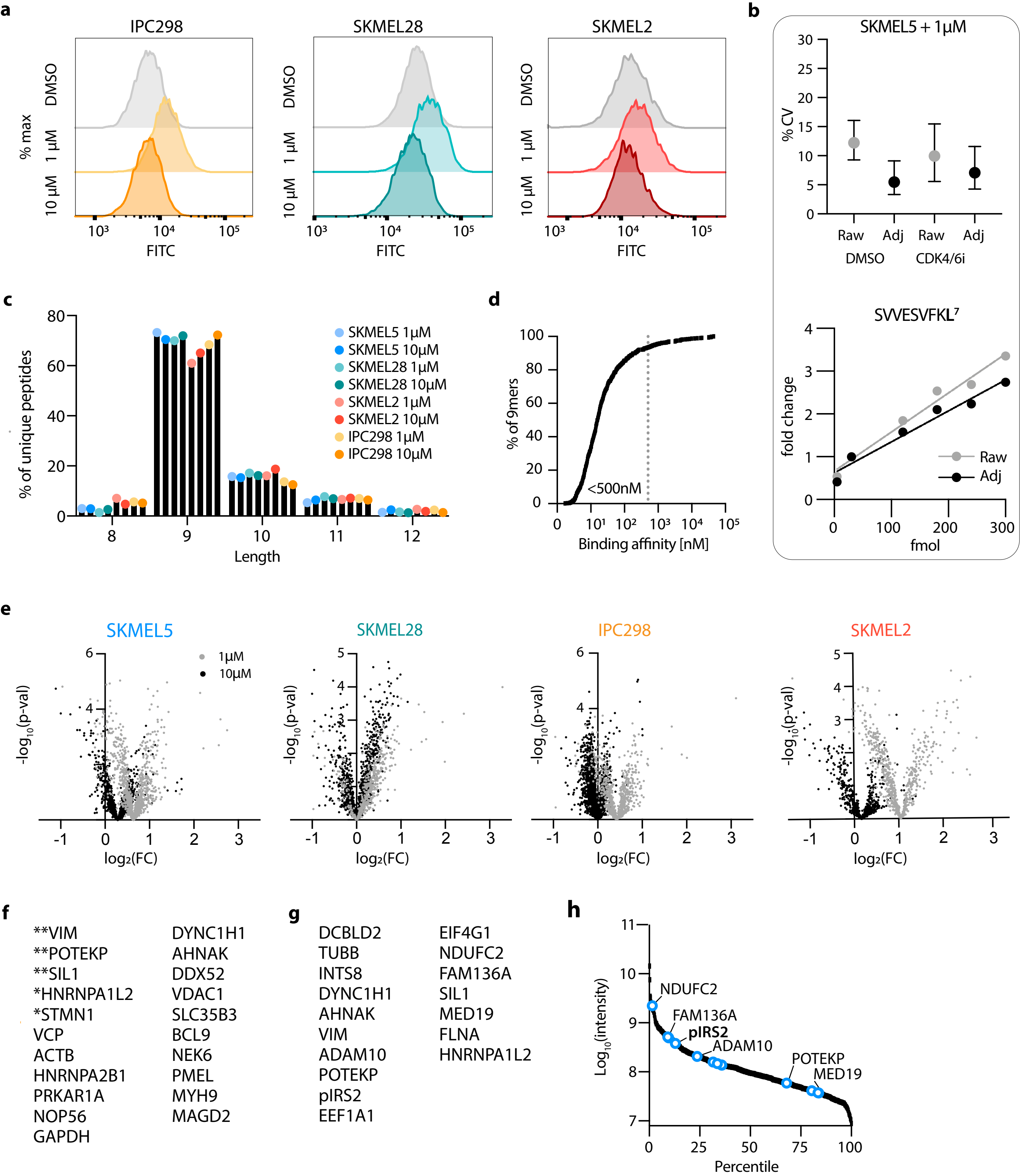

### Supplemental figure 4

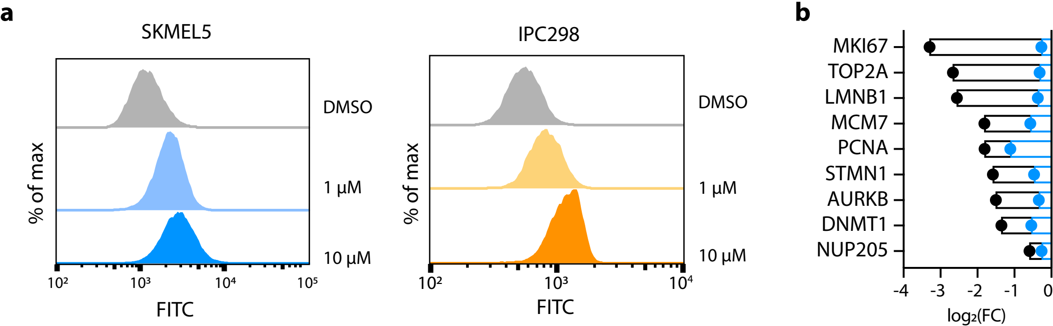

Supplementary figure 5

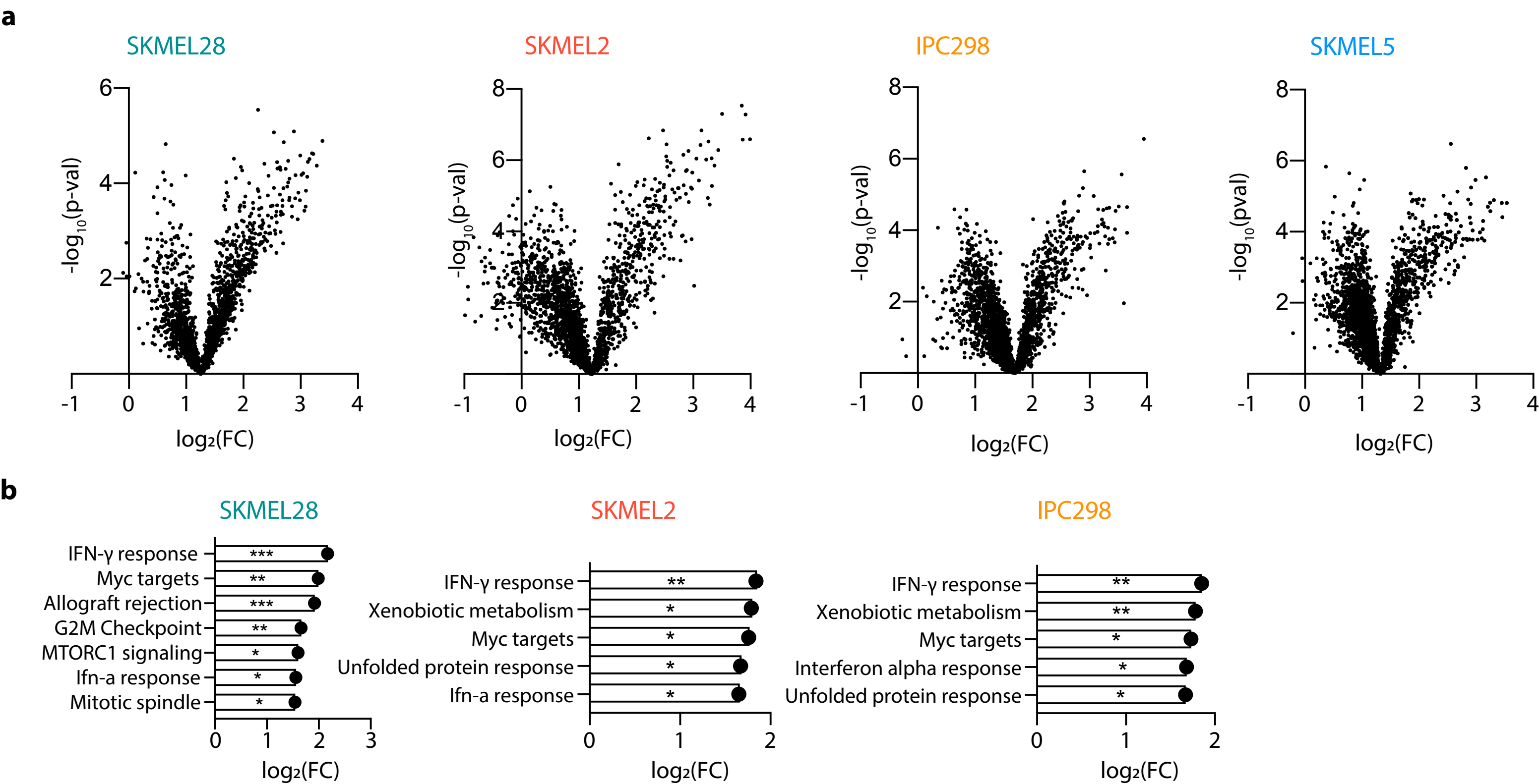

SUPPLEMENTARY FIGURE LEGENDS

**Supplementary Fig. 1. a**. LF analyses of MDA-MB-231 cells show poor overlap (orange) of identified peptides across five analyses compared to the number of peptides identified in a single analysis (black). **b.** Multiple analyses of MDA-MB-231 cells analyzing 15-20% of peptide elution per analysis show TMT-labeled samples begins to saturate unique peptides identified after four replicate analyses. **c.** Length distribution of MDA-MB-231 peptides for LF and TMT-labeled samples. **d.** Number of unique peptides identified in a comparison analysis using $1x10^7$ MDA-MB-231 cells for immunopeptidomic profiling vs. $5x10^6$ cells. Numbers represent a single analysis using 20% of the peptide elution.

**Supplementary Fig. 2.** Calibration curve of the DDX5 peptide (KLDVGNAEV) added into 1x107 MDA-MB-231 cells from 0.1 fmol to 1 fmol. 0.1 fmol corresponds to ~6 copies per cell.

**Supplementary Fig. 3. a.** Surface HLA expression measured via flow cytometry presented as % of maximum signal. A representative plot of distributions observed among three replicates is shown. **b.** Applying hipMHC correction factors decreases the coefficient of variation in both DMSO and 1 µM Palbociclib SKMEL5 cells (upper), and the titrated peptide, SVVESVFKL, displays 3.6x dynamic range suppression (lower). **c.** Length distribution of peptides identified in each cell line and treatment. Data is represented as % of total peptides identified. **d.** Predicted binding affinity of 9mer peptides in SKMEL5 cells with DMSO or 1 µM CDK4/6i. 93.3% have a predicted affinity of <500nM. **e.** Volcano plots representing $log_2$ fold change (FC) of CDK4/6i over DMSO on x-axis, and significance on y-axis, 1 µM (grey) and 10 µM (black). Significance (paired t test) for low dose treatment of SKMEL5, IPC298, and SKMEL2 cells was calculated after mean centering the treated data. **f.** Source proteins of peptides significantly enriched (mean adj) following CDK4/6i. **= significantly upregulated in four cell lines, *= three lines, all others were seen in at least two cell lines. **g.** 17 common source proteins significantly increasing in all four cell lines. **h.** $Log_{10}$ of average peptide AUC integrated abundance vs. percentile rank of all peptides. Blue points are peptides labeled by their source protein that are significantly increased in all four cell lines.

**Supplementary Figure 4. a.** Histogram of γ-H2AX levels determined by flow cytometry. Data is represented as % of maximum signal. **b.** $Log_2$(FC) for SKMEL5 E2F peptides significantly decreasing (blue) with 10µM CDK4/6i for 72h and matched RNA expression of SKMEL5 cells treated with 1µM CDK4/6i for 72h (black), as RNA sequencing was not performed on SKMEL5 cells treated with 10µM CDK4/6i.

**Supplementary Fig. 5. a**. Volcano plots of the IFN-γ regulated peptides. Displayed are the log2 fold change (FC) of IFN-γ stimulated abundances over DMSO versus significance (mean adjusted p-value). **b**. Enrichment analysis of IFN-γ stimulated cells. Plots represent normalized enrichment score (NES) of significant positively enriched Hallmark gene set pathways. No pathways were negatively enriched. $q < 0.25$, *$p < 0.05$, **$p < 0.01$, ***$p < 0.001$. **c.** Venn diagram displaying the number of unique peptides observed in SKMEL5 cells treated with 1 $\mu$M CDK4/6i and/or IFN-γ stimulation.
